# Laboratory diagnostics of tomato spotted wilt virus by PCR

**DOI:** 10.1101/2021.01.17.426990

**Authors:** O.N. Morozova, D. D. Zvyagintseva, O.O. Beloshapkina

**Affiliations:** All-Russian Science-Research Institute of Plant Quarantine; doctor of agricultural sciences, professor, Russian State Agrarian University - Moscow Timiryazev Agricultural Academy

**Keywords:** PCR, TSWV, tospoviruses, diagnostic of viruses

## Abstract

Tomato spotted wilt virus is a widespread and harmful virus. It affects a wide range of host plants. The outward signs of TSWV lesions are different on different cultures. For the early diagnosis of TSWV, molecular diagnostic methods such as PCR must be used. In the course of this work, primers were developed for the diagnosis of tomato bronzing virus by real-time PCR and classical PCR. We also compared the specificity and sensitivity of various test systems for the diagnosis of tomato bronzing virus. In the course of this comparison, it was found that the self-assembly test system and the Syntol test system can be recommended for laboratory diagnostics of tomato spotted wilt virus.

## Summary

Tomato spotted wilt virus (TSWV) is the typical member of genus Tospovirus, family Peribunyaviridae. It is included in List of non-quarantine harmful organisms, subjected to regulation on territory of Russian Federation, and in List A2 of EPPO. TSWV infects many vegetable and flower-ornamental crops and seriously reduces it’s productivity, decorativeness and vitality. The main hosts are peanut, brovallia, pelargonium, anemone, aster, balsam, begonia, gerbera, calceolaria, tagetes, cineraria, cyclamen, zinnia, chrysanthemum, tobacco, pepper, salad, tomato, eggplant, legumes and some other crops. According to various estimates, up to 900 plants from 90 families may be hosts of TSWV [1,2,3,4].

External signes of infection by TSWV are very variable at different crops. The main symptoms are chlorotic and necrotic ringspots, mottled leaves, the appearance of line patterns and yellow mesh on them, deformation of leaf blades and yellowing of vines, falling flowers, streak and mosaic of petals and fruits, systematic wilting of plant [5, 6].

Sucking insects - thrips are playing the main role in distribution of TSWV in nature and agrocenosises. The most important representors of this group are *Thrips tabaci, Frankliniella intonsa, Frankliniella occidentalis* [7, 8, 9, 10]. The last is quarantine pest, which is dangerous to flower and vegetable crops in greenhouses in Russia.

Currently, there has been a progressing dynamics of the spread of the tomato bronzing virus in many countries, incl. and in Russia. This virus, as well as its vectors, thrips, already have a stable circulation in natural foci in the south of Russia, as well as on various flower crops and tomatoes in open and protected ground. One of the most important measures to prevent the widespread acclimatization of the tomato spotted wilt virus is its early diagnosis in host plants using modern virological methods.

**The purpose** of this work was to refine and improve the elements of existing molecular methods for diagnosing tomato spotted wilt virus.

## Materials and methods

The work was carried out in the laboratory of virology and analysis of GMO ILC of All-Russian Science-Research Institute of Plant Quarantine. The testing of existing and developed methods of diagnostics of tospoviruses was carried out with isolates and positive controls of the following viruses:

1. Impatiens necrotic spot virus (INSV) – isolate PV-0280 (DSMZ, Germany).
2. Tomato spotted wilt virus (TSWV) – isolates PV-0182, PV-0204, PV-0393 (DSMZ, Germany).
3. Tomato spotted wilt virus (TSWV) – positive control (LOEWE, Germany).
4. Watermelon silver mottle virus (WSMoV) – isolate PV-0283 (DSMZ, Germany).
5. Iris Yellow Spot Virus (IYSV) – isolate PV-0528 (DSMZ, Germany).
6. Chrysantemum stem necrosis virus (CSNV) – isolate PV-0529 (DSMZ, Germany).
7. Negative control (tobacco leaves) (Agdia, USA)

When working out the test based on PCR, the above-mentioned lyophilized reference isolates from DSMZ (Germany), positive and negative controls for enzyme-linked immunosorbent assay, potted plants of ornamental crops, and healthy plants were used.

RNA isolation was performed with reagent kits for nucleic acid isolation (Proba-NK) (Agrodiagnostika) and FITO-SORB (Syntol) (both Russia), according to the manufacturer’s protocol. The reverse transcription reaction was performed using reagents for RT with an OT-Random primer (Agrodiagnostika) and a kit of reagents MMLVRTkit (Evrogen, Russia), according to the manufacturer’s protocol. For the amplification reaction, a set of PCR reagents Master-mix ScreenMix HS (Evrogen, Russia), 5xMasCFGTaqMIX-2025 (unstained) and 5xMasDDTaqMIX-2025 (stained) (Dialat, Russia) was used in accordance with the manufacturer’s protocol. PCR was carried out on amplifiers CFX - 96, C1000 (BioRad, USA), DT-Prime (DNA-Technologies, Russia).

The following primers and a probe were used for real-time PCR (synthesis at Evrogen, Russia):

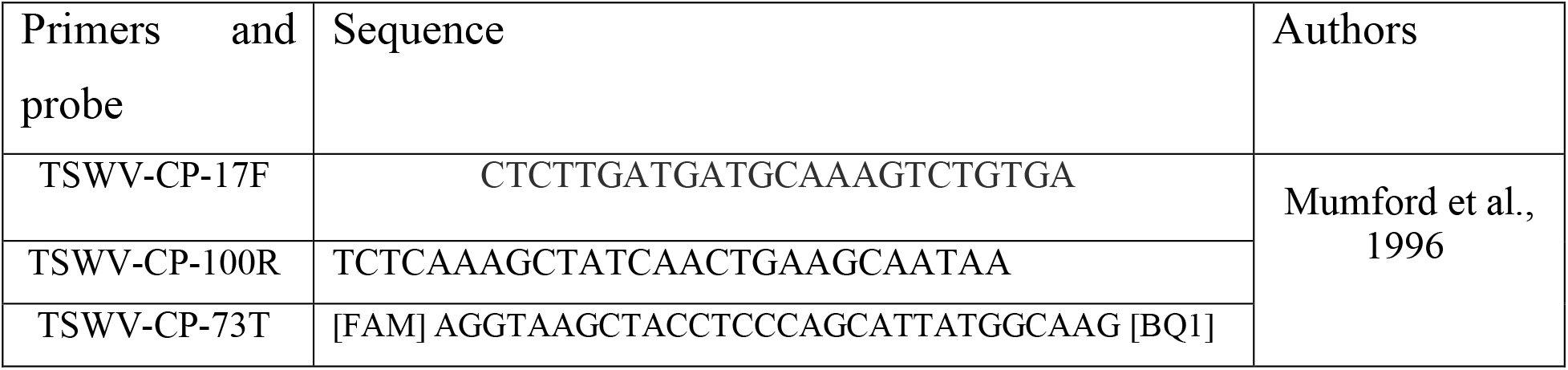

The conditions for the amplification with the TSWV-CP-17F / TSWV-CP-100R soligonucleotides, the TSWV-CP-73T probe are as follows: 5 min at 95 ° C; reaction within 45 cycles: 15 sec at 95 ° C, 45 sec at 55 ° C; 4 ° C ∞. Species-specific probe TSWV-CP-73T allows detecting a specific fragment for Tomato spotted wilt virus in the reaction mixture via the FAM / Green fluorescence channel.

For comparison with the self-assembly test system, commercial TSWV detection kits from Agrodiagnostika (Russia) and Syntol (Russia) were used.

The following primers were used for classical PCR:

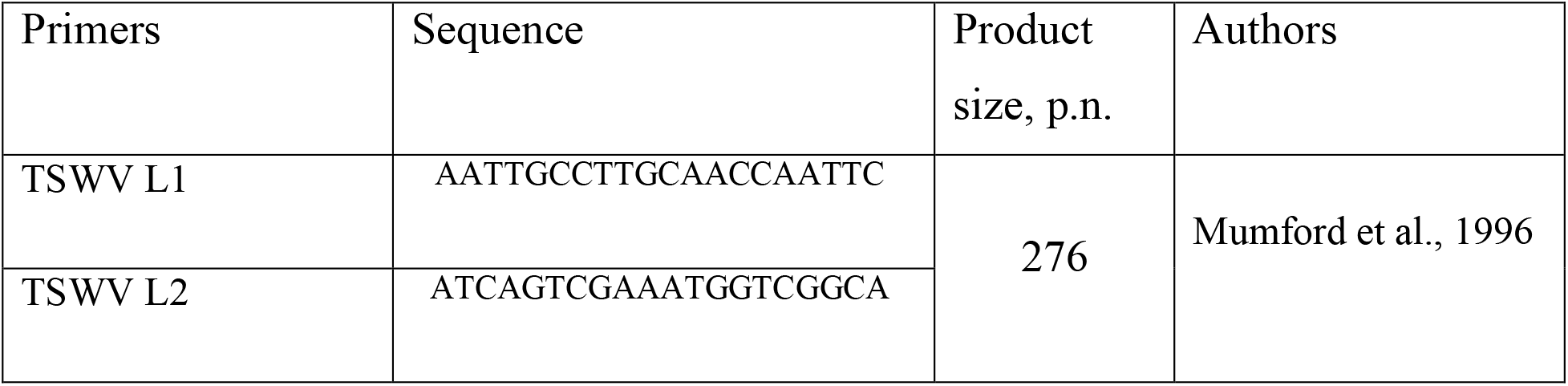

The conditions for carrying out amplification with TSWV L1 / TSWV L2 oligonucleotides are as follows: 10 min at 96 ° C, reaction for 40 cycles: 60 sec at 94 ° C, 60 sec at 55 ° C, 60 sec at 72 ° C; 10 min at 72 ° C, 4 ° C ∞. The detection of the results of classical RT-PCR was carried out by electrophoresis in 1.5% agarose gel.

## Results and discussion

In the course of this work, specific primers TSWV-CP-17F / TSWV-CP-100R and a species-specific probe TSWV-CP-73T (synthesis at Evrogen, Russia) were tested to identify the causative agent of tomato spotted wilt vitus from plant material. Amplification was performed in real time with positive control samples. Figure 2 shows the dependence of the fluorescence of the FAM / Green channel on the cycle number. According to the results of the amplification reaction in Figure 2, it can be seen that only positive controls of the tomato bronzing virus worked. PKO + 1 (TSWV PV-0182) detected at 23.47 Ct, PKO + 2 (TSWV PV-0182) at 22.9 Ct, PKO + 3 (TSWV PV-0182) at 19.08 Ct, where Ct - the number of the threshold cycle.

**Figure 1.**
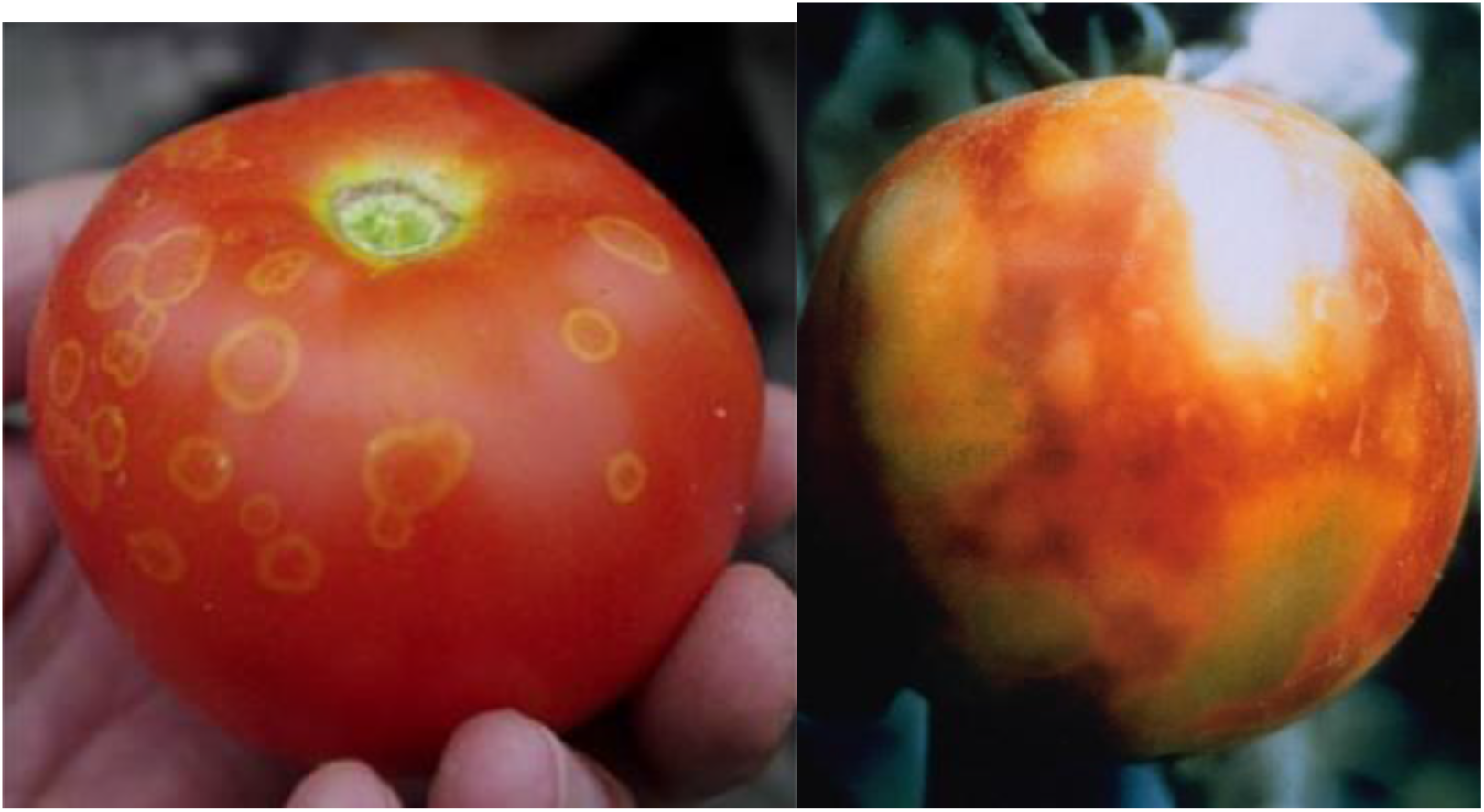
Symptoms of TSWV on tomatoes. (photo by G. Marchoux, INRA, France and P. Mariman, USA) [4]

**Figure 2.**
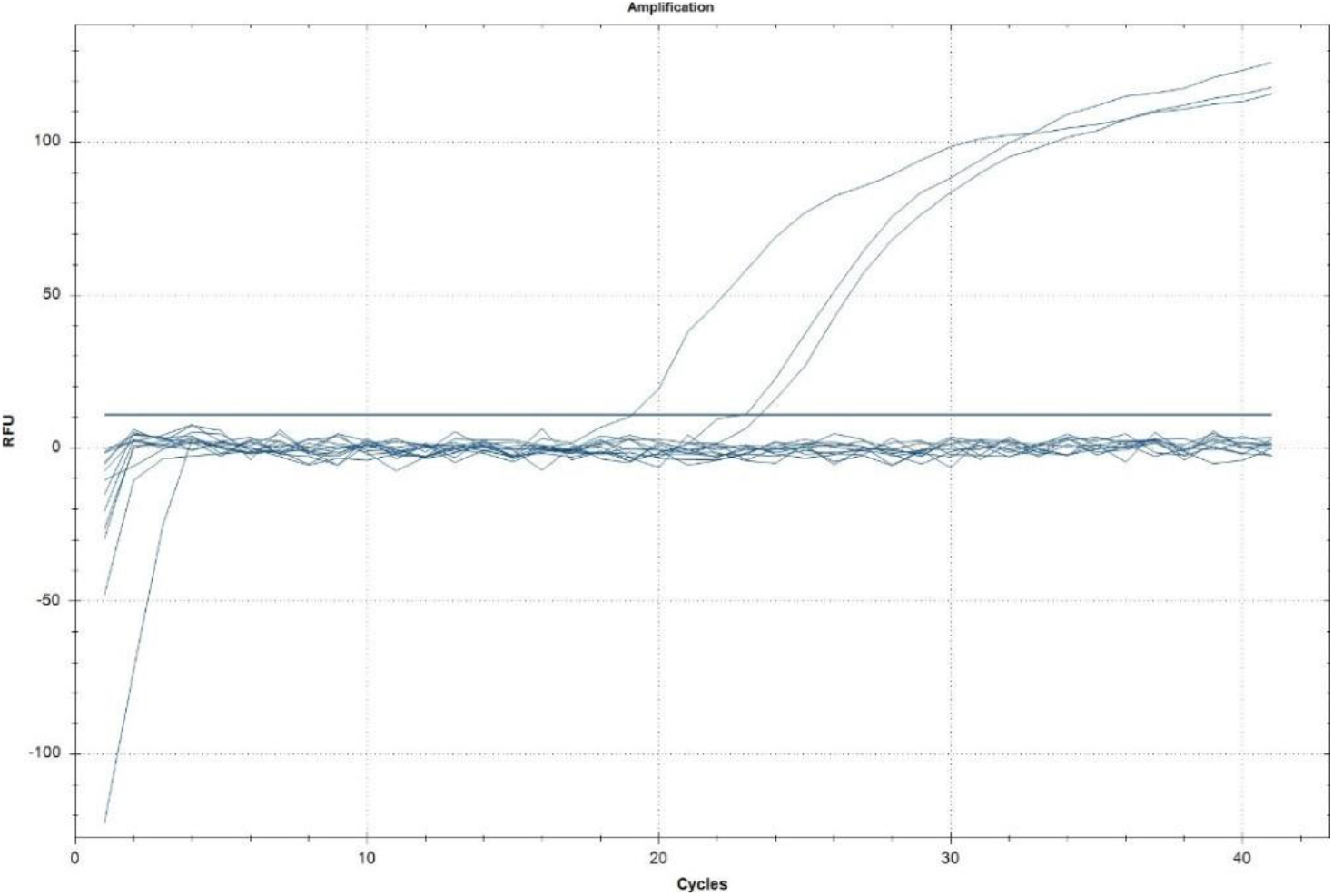
Results of PCR in real time: dependence of the fluorescence of the FAM / Green channel (TSWV)

Amplification of cDNA of the causative agent of tomato bronzing with the same samples (Fig. 3) with a pair of primers TSWV L1 / TSWV L2 with a ready-made mixture 5x MasCFEMIX-2025 (Dialat, Russia) showed that the studied samples with cDNA of the virus had the desired amplification product of a specific size 276 p.n. At the same time, there was no nonspecific reaction with other closely related species of tospoviruses.

**Figure 3.**
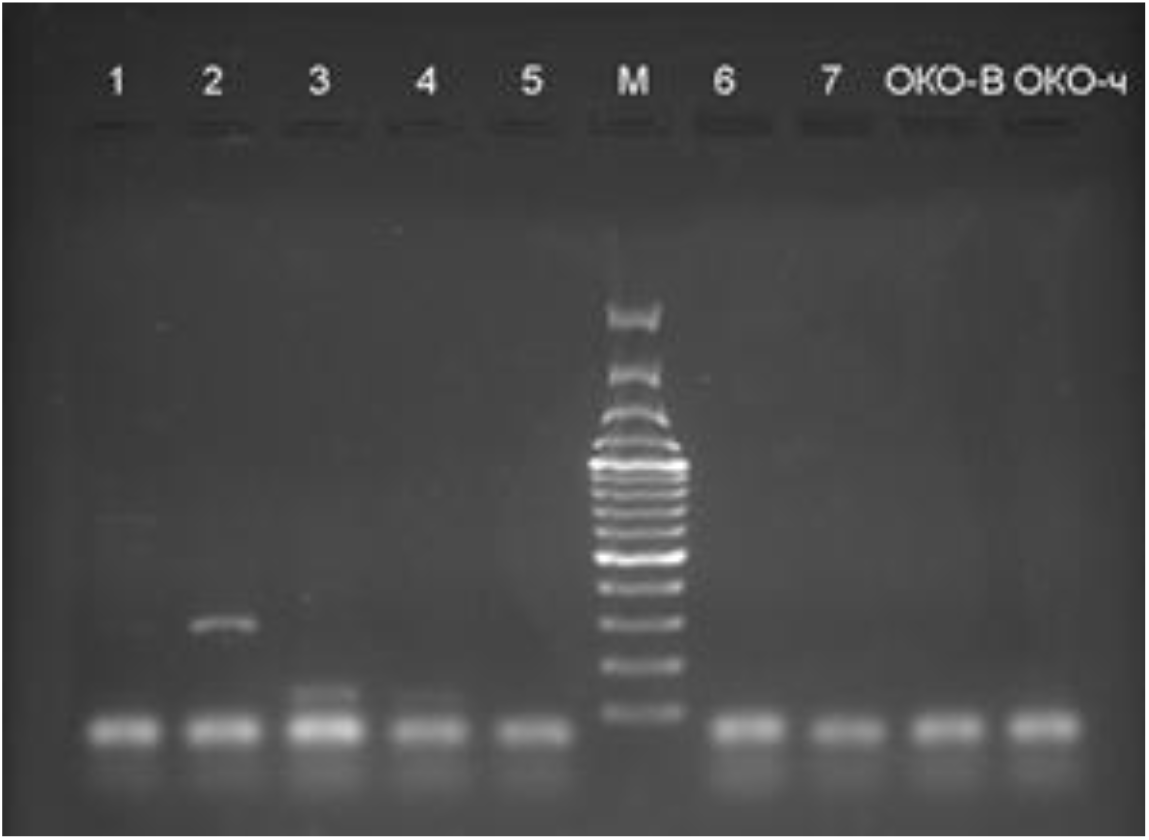
Electropherogram of classical PCR samples with TSWV L1 / TSWV L2 primers. Where: 1 - isolate TSWV PV-0182; 2 - isolate TSWV PV-0204; 3 - chrysanthemum plant; 4 - chrysanthemum plant; 5 - isolate INSV PV-0280; 6 - isolate WSMoV PV-0283; 7 - CSNV PV-0529; OKO-V - negative control of secretion; OKO-h - negative control of the clear zone. M - molecular weight marker 100-3000 bp. (“Thermo Fisher scientific”, USA).

It can be concluded that the classical PCR method using the TSWV L1 / TSWV L2 primer pair is specific for the detection of tomato spotted wilt virus. Although the sensitivity of this method is slightly lower than real-time PCR, it is quite satisfactory and provides reliable results.

Table 1 shows a comparative assessment of the results of real-time PCR for various test systems.

**Table 1.** Comparative assessment of tomato spotted wilt virus detection by PCR with different test systems in real time

| Sample | Test system of the firm<br>"Agrodiagnostika" |  | Test system of the firm<br>"Syntol" |  | Test system with primers<br>" TSWV-CP-17F/ TSWV -CP-100R и<br>зондом TSWV -CP-73T |  |
| --- | --- | --- | --- | --- | --- | --- |
|  | Result | Threshold<br>cycle. (Ct) | Result | Threshold<br>cycle. (Ct) | Result | Threshold<br>cycle. (Ct) |
| IYSV_<br>PV-0528 | - | - | - | - | - | - |
| TSWV_L<br>oewe | - | - | + | **20,2** | + | **21,7** |
| TSWV_P<br>V-0393 | + | **31,27** | + | **19,6** | + | **23,1** |
| TSWV_<br>PV-0204 | - | - | + | **19,9** | + | **18,8** |
| INSV_PV-0280 | - | - | - | - | - | - |
| TSWV_<br>PV-0282 | + | **29,83** | + | **17,9** | + | **20,8** |
| TSWV_<br>PV-0282_1/10 | + | **33,12** | + | **20,9** | + | **24,3** |
| TSWV_<br>PV-0282_1/10<br>0 | + | **36,99** | + | **24,9** | + | **27,4** |
| TSWV_<br>PV-0282_1/10<br>00 | + | **40,76** | + | **28,5** | + | **31,2** |
| TSWV_P<br>V-0282_1/10<br>000 | - | - | + | **33,4** | + | **35,7** |
| TSWV_P<br>V-0282_1/10<br>0000 | - | - | + | **36,3** | + | **37,5** |
| CSNV PV-0529 | - | - | - | - | - | - |
| WSMoV P<br>V-0283 | - | - | - | - | - | - |
| Negative<br>control | - | - | - | - | - | - |

Based on our data, it can be argued that the Agrodiagnostika test system is the least specific and sensitive. Two of the samples tested gave a false negative result; and when the sample was diluted 10,000 times, the test result was negative. At the same time, the same samples gave a positive result when tested with a kit from Syntol, as well as a self-assembly system with primers TSWV-CP-17F / TSWV-CP-100R and a species-specific probe TSWV-CP-73T.

It should be noted that none of the tested test systems showed a false positive result when studying isolates of related tospoviruses.

## Conclusion

Tomato spotted wilt tospovirus has a wide range of host plants among economically significant vegetable and flower-ornamental crops, causing significant damage to them all over the world. Effective protection against it can be organized only on the basis of regular reliable phytosanitary monitoring, which is impossible without the use of modern highly sensitive methods of laboratory examination. In the course of the study, we have worked out test systems for PCR both in real time and classical with detection by electrophoresis. The specificity and sensitivity of three different real-time PCR test systems was evaluated. It has been proven that a set of reagents from Syntol (Russia) and a self-assembly system with primers TSWV-CP-17F / TSWV-CP-100R and a species-specific probe TSWV-CP-73T can be recommended for laboratory diagnostics of tomato spotted wilt virus.

## References

1. Scholthof K-B.G., Adkins S., Czosnek H., Palukaitis P., Jacquot E., Hohn T., Hohn B., Saunders K., Candresse T., Ahlquist P., Hemenway C., Foster G.D. Top 10 plant viruses in molecular plant pathology // Molecular Plant Pathology. – 2011. – Vol. 12. – P. 938–954.

2. CABI, 2020. CABI Crop Protection Compendium // CAB International, Wallingford, UK.

3. EFSA, 2012. Scientific opinion on the pest categorization of the tospoviruses // EFSA Panel on Plant Health/EFSA Journal. – 2012. – Vol. 10. – P. 1–32.

4. EPPO Global Database, 2020. http://www.eppo.int/DATABASES/pqr/pqr.htm.

5. EPPO, 2004. Diagnostic protocol for regulated pests. PM 7/34(1). Tomato spotted wilt tospovirus, Impatiens necrotic spot tospovirus and Watermelon silver mottle tospovirus // Bulletin OEPP/EPPO Bulletin. – 2004. – Vol. 34. – P. 271–279.

6. Pappu H.R., Jones R.A.C., Jain R.K. Global status of tospovirus epidemics in diverse cropping systems: Successes achieved and challenges ahead // Virus Research. – 2009. – Vol. 141. – P. 219–236.

7. EFSA, 2012. Scientific opinion on the pest categorization of the tospoviruses // EFSA Panel on Plant Health/EFSA Journal. – 2012. – Vol. 10. – P. 1–32.

8. Avila Y., Stavisky J., Hague S., Funderburk J., Reitz S., Momol T. Evaluation of *Frankliniella bispinosa (Thysanoptera : Thripidae*) as a vector of the *Tomato spotted wilt virus* in pepper // Florida Entomologist. – 2006. – Vol. 89. – P. 204–207.

9. de Borbon C.M., Gracia O., Piccolo R. Relationships between tospovirus incidence and thrips populations on tomato in Mendoza, Argentina // Journal of Phytopathology. – 2006. – Vol. 154. – P. 93–99.

10. Fujisawa I., Tanaka K., Ishii M. Tomato spotted wilt virus transmissibility by three species of thrips, *Thrips setosus*, *Thrips tabaci* and *Thrips palmi* // Annals of the Phytopathological Society of Japan. – 1988. – Vol. 54. – P. 392.

11. Mumford R.A., Barker I., Wood K.R. An improved method for the detection of Tospoviruses using the polymerase chain reaction // Journal of Virological Methods. – 1996b. – Vol. 57. – P. 109–115

